# Tracing the drivers of range-wide bowhead whale genomic structure and diversity

**DOI:** 10.1101/2025.05.23.655713

**Authors:** Michael V. Westbury, Nicholas A. Freymueller, Andrea A. Cabrera, Lutz Bachmann, Steven H. Ferguson, Mads Peter Heide-Jørgensen, Kit M. Kovacs, Christian Lydersen, Olga Shpak, Øystein Wiig, Damien A. Fordham, Eline D. Lorenzen

## Abstract

Understanding how past environmental and anthropogenic factors shaped contemporary genetic patterns is essential for assessing the vulnerability of species in the face of ongoing climate change. The bowhead whale (*Balaena mysticetus*) – the only baleen whale found in the Arctic year-round – has a strong association with sea ice, long history of human exploitation, and a circumpolar distribution, making it a valuable model for investigating how these factors have shaped population structure in the marine Arctic. We analysed both nuclear and mitochondrial genomes from bowhead whale individuals sampled across the species’ circumpolar range, encompassing all four recognized management stocks. Our results reveal three not four genetically differentiated populations: the Sea of Okhotsk stock; the East Greenland-Svalbard-Barents Sea stock; and a combined population containing the Bering-Chukchi-Beaufort Seas and East Canada-West Greenland stocks. We utilised ecological niche modeling and bowhead whale telemetry data to model stock connectivity over the past 11,700 years, and identify past habitat connectivity as a key driver of current population structure. Despite centuries of intensive commercial whaling, the three bowhead whale populations exhibit little-to-no evidence of recent inbreeding and retain high genetic diversity relative to other mammalian species. The lowest genetic diversity, most inbreeding, and highest realised genetic load being in the Sea of Okhotsk population. Collectively, our findings shed light on the recent population history and dynamics of bowhead whales, and offer valuable baseline data on present-day genetic structure and diversity to support effective conservation and management strategies.

## Introduction

Interactions between genetic and ecological processes are fundamental to understanding contemporary patterns of phylogeographic structure and diversity and how they have been impacted by global change^1^. This is because contemporary genetic variation is conditioned by past climatic and environmental change along with anthropogenic influences that individually, or in concert, drive population structure and adaptive potential ^2^. Understanding their relative roles is essential for determining current and future threats to the long-term survival of a species.

In marine ecosystems, anthropogenic drivers have modified species’ contemporary genetic structure and diversity differently than in terrestrial systems. The latter have been subject to human-induced pressures across many millennia, including through direct hunting, changes in fire regimes, and landscape modification ^3^. In contrast, marine environments have remained largely unaffected by anthropogenic influences until the past ∼500 years ^4^. Consequently, contemporary genetic patterns in marine species are inferred to be more directly linked to natural environmental drivers ^5^, although anthropogenic impacts, such as ongoing climate change and industrial exploitation, are becoming more pronounced ^4^.

Bowhead whales (*Balaena mysticetus*) are the only baleen whale endemic to the Arctic. Populations of bowhead whales suffered major declines across their circumpolar range during four centuries of commercial whaling (∼1530-1914) ^6^. They were ideal targets due to their slow swimming speed, relatively docile nature, and ease of retrieval after killing. International agreements giving bowhead whales formal protection did not come into effect until 1931. That is after commercial exploitation had already been abandoned due to whaling no longer being profitabile ^7^.

Bowhead whales live along the loose pack edges of Arctic sea ice, where they feed on large quantities of zooplankton. Their annual movement patterns are primarily governed by changes in sea ice cover ^8^. The fossil record for bowhead whales, which is rich compared to other marine mammals ^8,9^, indicates that distributional shifts during the Late Quaternary resulted from shifting sea ice conditions ^9,10^. However, palaeogenomics and habitat modelling of bowhead whales in the Atlantic sector of the Arctic suggest population stability across the Holocene (past 11,700 years), despite environmental perturbations ^11^. At a global scale, suitable bowhead whale habitat inferred from ecological niche models is forecast to rapidly decline by the end of the 21st century, indicating an uncertain future for this charismatic species ^12,13^.

The International Whaling Commission (IWC), which sets overarching conservation guidelines of bowhead whales across their circumpolar range, currently recognises four geographically segregated management stocks (Fig. 1A), based on satellite telemetry and genetic data ^14^. These stocks are named: Bering-Chukchi-Beaufort Seas (BCB); East Canada-West Greenland (ECWG), which was previously split into the Hudson Bay and David Strait stocks ^15^; East Greenland-Svalbard-Barents Sea (EGSB); and the Sea of Okhotsk (OKH) ^16^. Bowhead whales undertake seasonal movements within these stock boundaries, and rare dispersal events between different stocks have only recently been detected ^17^. Whether this observed connectivity will persist in the future and whether it has occurred regularly in the past remains unknown.

**Figure 1:**
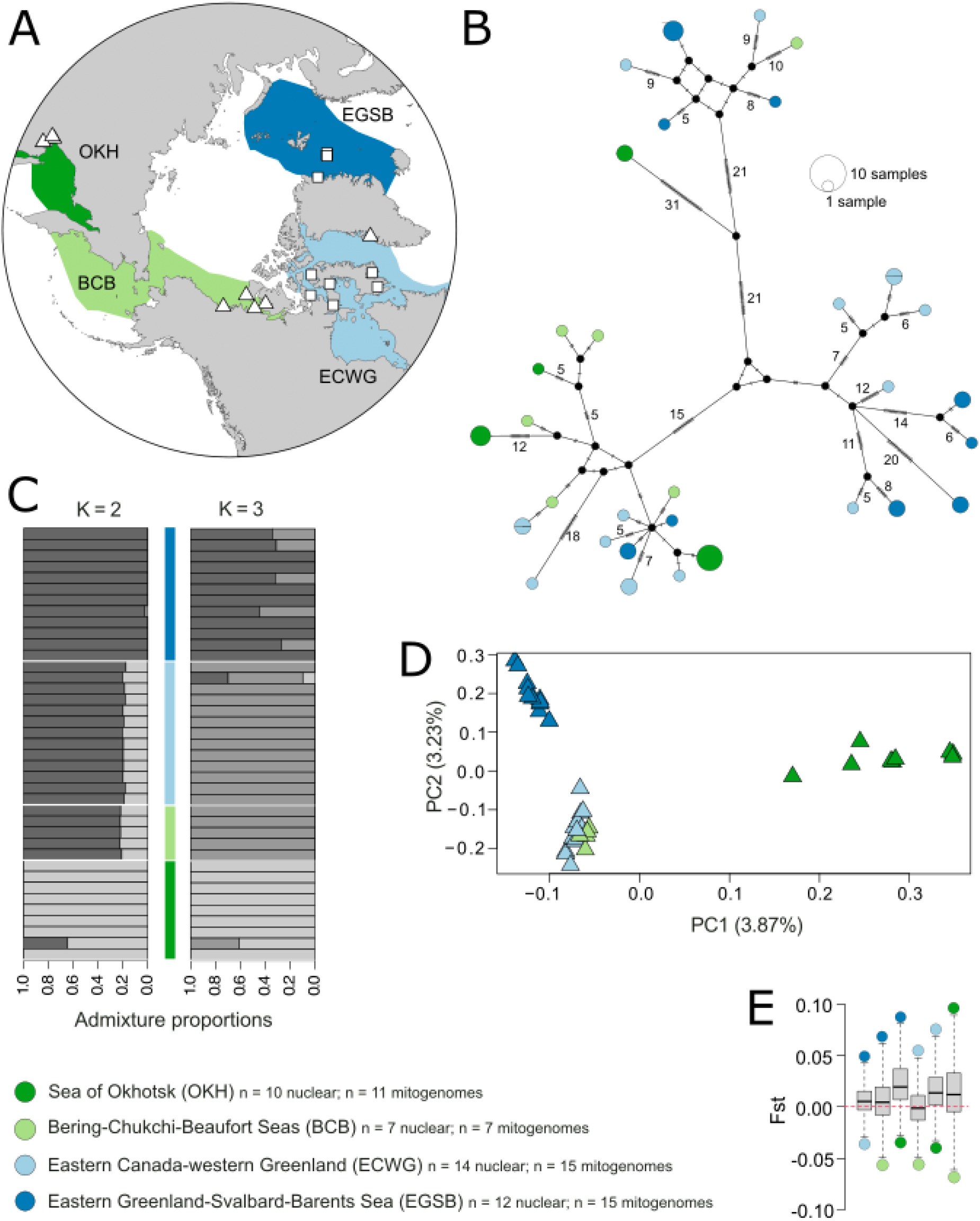
Genetic population structure among stocks of bowhead whales. **A)** Map of sample localities (white symbols; triangles indicate the 24 samples new to this study and squares denote existing samples) and the four bowhead whale management stock boundaries which bowhead whales regularly travel throughout during the year (coloured shading). **B)** Median-joining haplotype network constructed in PopART, based on the 32 haplotypes present among 48 mitochondrial genomes. Haplotypes are represented by circles, the relative size of which are proportional to the number of individuals. Colours represent stock of origin. Missing haplotypes are indicated by black dots. Hashes indicate the number of substitutions among haplotypes, denoted by a number for > 5. **C)** Nuclear genomic admixture proportions created using genotype likelihoods and number of populations set at K = 2 and K = 3. **D)** Nuclear genomic Principle Components Analysis (PCA) created from pseudohaploid base calls. Percentages of variation explained by PC1 and PC2 are indicated in parentheses. **E)** Nuclear genomic *F*_ST_ values between individuals, grouped into the four bowhead whale management stocks.

Patterns of genetic structure and diversity across the circumpolar range of bowhead whales are not fully understood. Previous studies have used genetic markers with limited power to discern structure, or have had a regional focus ^14^. Maternally inherited short mtDNA regions have been used to uncover population size changes over time and a lack of local population structure ^18–20^. Microsatellite studies found some evidence of population structure within the BCB stock ^21^, but this was later refuted ^22^ due to inherent challenges with reproducibility and reliability of microsatellite data. More recently, whole-genome sequencing has led to more robust and reproducible stock-specific demographic analyses ^11,23–25^. However, these efforts have focused on individual stocks or specific geographic regions, leaving significant gaps in our understanding of genetic structure from a range-wide perspective. Comprehensive genomic analyses across the full distribution range of the bowhead whales are necessary to provide a holistic view of their genetic diversity, population structure and connectivity, and demographic history.

Past environmental factors influencing bowhead whale distribution have been explored through ecological niche models, offering reconstructions of past and future habitat suitability ^10–12^. These studies have primarily focused on quantifying changes in suitable habitable area over the Holocene and forecasting potential changes in coming decades. However, they are yet to be used to explore the spatial structure of suitable habitat patches through time, or to quantify genetic connectivity patterns between different bowhead whale stocks. Doing so could provide important insights into the role of climate and environment in shaping current genetic structure and diversity.

Here, we explore determinants of range-wide patterns of contemporary genomic population structure and diversity, and past demographic history for bowhead whales, to establish an ecological and evolutionary baseline for understanding the threats from future climatic change and increased human activity in the Arctic. We analyse nuclear genomes from 39 contemporary bowhead whale individuals and 48 mitochondrial genomes sampled across the four management stocks (Fig. 1A). We evaluate whether changing environments shaped bowhead whale phylogeography and range-wide population structure by interpreting our genomic findings alongside estimates of inter-stock connectivity across the Holocene, derived from bowhead whale ecological niche models ^13^ and movement data ^12,26^.

## Results

### Genomic population structure

To investigate population subdivision and connectivity from a nuclear genomic perspective, we conducted principal component analysis (PCA), a neighbour-joining tree, admixture proportion estimates, fixation index (*F*_ST_) calculations, and D-statistics. To evaluate maternal genetic structure, we conducted a haplotype network analysis and *F*_ST_ calculations based on the complete mitochondrial genomes.

Regardless of the base call approach of the nuclear data (genotype likelihoods or pseudohaploid), bowhead whales from OKH separated from the other stocks along the PC1 axis (Fig. 1D and Supplementary Figure S5). When using pseudohaploids, EGSB individuals separated from the BCB and ECWG along the PC2 axis, regardless of whether EGSB individuals were downsampled to 2x or not (Fig 1D and Supplementary Figure S6A). The EGSB individuals were widely spread along PC2 when using genotype likelihoods (Supplementary Figure S5). However, when downsampling these EGSB individuals to 2x and using genotype likelihoods, they formed a more distinct cluster (Supplementary Figure S6B). Regardless of the choice of method, individuals from the BCB and ECWG overlapped heavily, and likely comprise a single cluster. We therefore pooled individuals from these two stocks together for downstream analyses and henceforth refer to them as the ‘BCB/ECWG population’. A Tracy-Wildon test on the eigenvalues showed that only PC1 was significant (p<0.01) regardless of the base call approach or whether EGSB individuals were downsampled to 2x. The neighbour-joining tree supported the PCA results, with EGSB individuals and OKH individuals each forming their own clades, and the combined BCB and ECWG individuals forming a clade (Supplementary Figure S7). Although none of the 32 haplotypes were shared among individuals across stocks, the mitochondrial haplotype network did not show any clear phylogeographic structuring (Fig. 1B) with individuals from each stock distributed across the network.

In the admixture proportions analysis, only K = 2 populations converged within 50 replicates. OKH contained mostly ancestry from one population, EGSB contained mostly ancestry from the other population, and BCB/ECWG individuals contained mixed ancestry, overlapping ∼20% with OKH individuals and ∼80% with EGSB individuals (Fig. 1C). Although it did not converge, we also looked into the replicate with the highest likelihood for K=3. OKH individuals contained mostly unique ancestry, BCB/ECWG individuals contained mostly unique ancestry from a second population, and EGSB contained mixed ancestry between the second population (predominant in BCB/ECWG) and its own unique ancestry (Fig. 1C).

The *F*_ST_ values show a similar pattern to the PCA, with OKH being the most differentiated, followed by EGSB (Fig. 1E). When considering median values, only the comparison between individuals from the BCB and ECWG stocks gave an *F*_ST_ < 0 (−0.0016091), suggesting no clear genetic separation of these stocks. However, the mean *F*_ST_ was > 0 (0.0007873205). This was consistent when downsampling ECWG individuals to 2x (Supplementary Fig. S8). In our mitochondrial analysis, the majority of *F*_ST_ comparisons were non-significant (Supplementary Table S2). Only OKH vs EGSB was significant at a threshold of 0.01.

Our D-statistics results were mostly non-significant (Z < |3|) (Supplementary Fig. S9). The D-scores from the topology ((EGSB, ECWG), OKH) deviated most from 0. Many were negative, indicating some ancestral gene flow between EGSB and OKH relative to ECWG. However, there were also highly positive values indicating the opposite pattern. A similar pattern is seen for the topology ((BCB, ECWG), OKH). The topology ((EGSB, BCB), OKH) returned almost exclusively non-significant values. The topology ((ECWG, BCB), EGSB) returned some significant positive and negative values.

### Genomic diversity

To evaluate the genomic diversity and health of bowhead whale populations, we applied a range of complementary metrics to analyse the nuclear genomes. These included individual autosomal heterozygosity, population-level nucleotide diversity across both nuclear and mitochondrial genomes, autosomal runs of homozygosity (ROH), and estimates of nuclear genomic load based on the relative prevalence of missense and loss-of-function mutations.

Based on the three high-coverage nuclear genomes chosen as representative of each of the three genetic clusters identified in the structure analyses, we find the lowest mean genome-wide heterozygosity in the OKH individual (0.000979), followed by the EGSB individual (0.001067) (Fig. 2A,B). The ECWG individual had the highest heterozygosity (0.001226). These values are high relative to other mammalian species (Fig. 2A). Visualising the distribution of heterozygosity across the genome from the ANGSD output showed relatively similar distributions across individuals, apart from the OKH individual (CGG_1_024474), which showed relatively high counts of windows showing low-to-no levels of heterozygosity (Fig. 2A).

**Figure 2:**
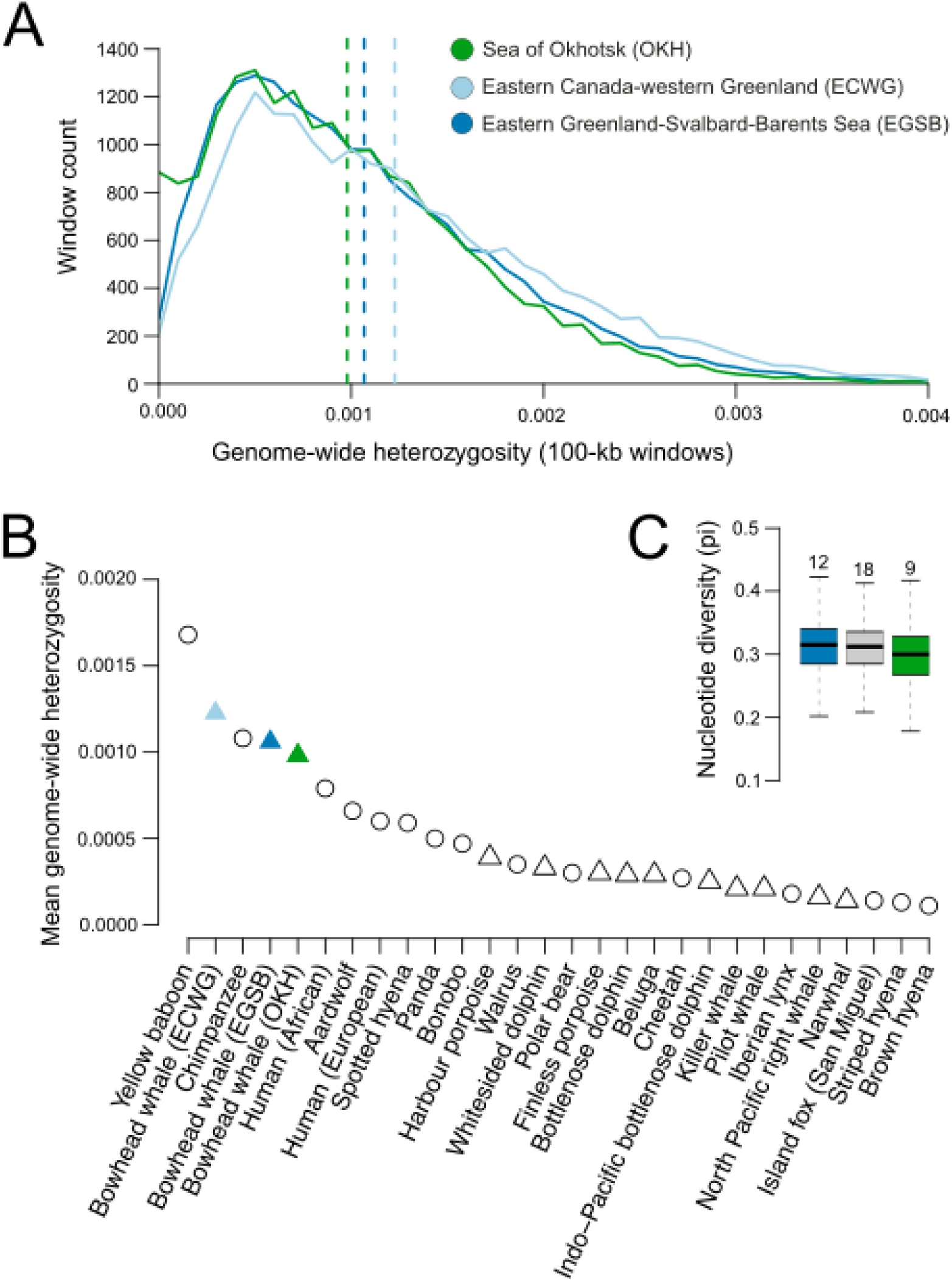
Genetic diversity of the three inferred bowhead whale populations. **A)** Nuclear genome-wide heterozygosity estimated for our three representative individuals, broken down into 100 kb windows; an individual from ECWG was used to represent the inferred combined BCB/ECWG population. Dotted lines indicate the mean values. **B)** Mean genome-wide heterozygosity of the three bowhead whale individuals and 25 other mammal species, taken from ^23^. Triangles indicate cetacean species. **C)** Population-level nuclear genome-wide nucleotide diversity calculated from pseudohaploid base calls in 100 kb windows. Here, individuals from the BCB and ECWG stocks were pooled and treated as one population (shown in grey) based on the structure analyses (Fig 1). OKH and EGSB were considered as separate populations. Sample size is indicated at the top of the range CV lines/bars.

When correcting for differences in sequencing coverage of the population-level dataset, OKH again had the lowest estimate of nuclear genomic nucleotide diversity (0.295). Similar levels of diversity were found in EGSB (0.310) and ECWG/BCB (0.308) (Fig. 2C).

In the 48 mitochondrial genomes analysed, we found high levels of haplotype and nucleotide diversity across the four stocks (Fig 1B, Supplementary Table S3). The OKH individuals showed the lowest haplotype and nucleotide diversity. ECWG had relatively low nucleotide diversity, but high haplotype diversity.

Estimates of inbreeding using ROHan only found runs of homozygosity > 1 Mb in the OKH individual, constituting 2.47% (1.65-2.73%) of the genome investigated and a mean length of 1,833,330 bp. No ROH >1 Mb were found in the other two individuals. The inbreeding coefficients from NGSrelate also did not show signs that any individual was inbred, with all F values being < 0.00012.

When considering genetic load due to both missense and/or loss-of-function mutations, EGSB had the highest masked genetic load, and ECWG had the lowest (Supplementary table S4). The highest realised load based on missense or loss of function mutations was found in OKH.

### Demographic history

To investigate deep-time demographic trends of the different bowhead whale populations, we implemented a pairwise sequential markovian coalescence model (PSMC) on the nuclear genomes of the three high coverage individuals. The three bowhead whale genomes show identical demographic trajectories of effective population size (N_e_) until ∼200 thousand years ago (kya), when the Okhotsk genome diverged from the other two (Fig 3A). At this time, the Okhotsk individual’s effective population size continues its trajectory of increasing N_e_, whereas the Canadian and Svalbard genomes plateau. At ∼120 kya, the two latter individuals decrease in N_e_ until ∼50 kya, after which they increase again. The Okhotsk N_e_ also begins to decrease ∼50 kya.

**Figure 3:**
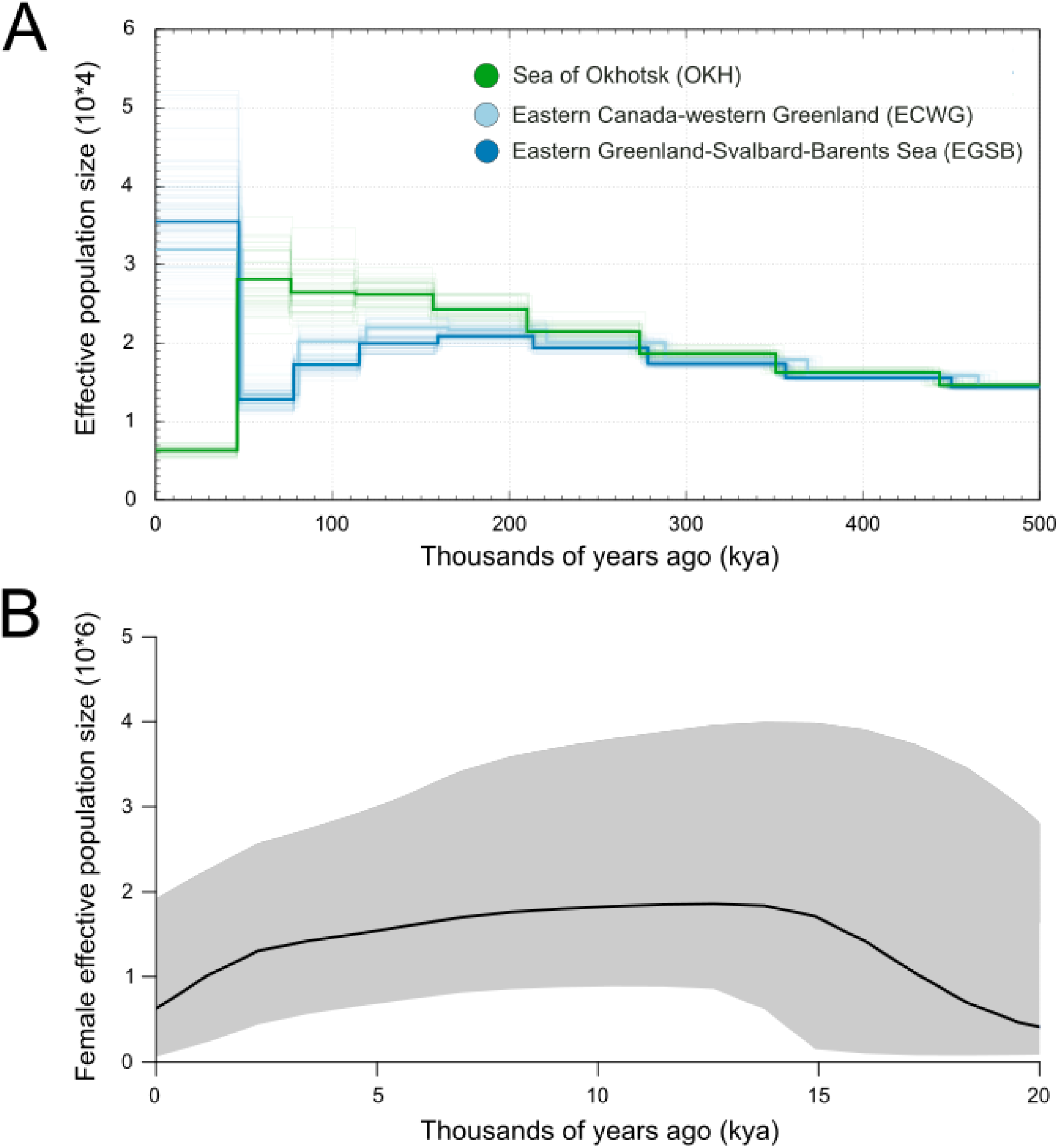
Bowhead whale demographic reconstruction. **A)** Deep-time demographic history calculated from our three representative individuals, using PSMC; an individual from ECWG was used to represent the inferred BCB/ECWG population. Faded lines indicated bootstrap replicates. **B)** More recent female demographic history based on a Skyline analysis of the mitochondrial genomes of 37 contemporary bowhead whale individuals, combined with 105 Holocene and 2 Late Pleistocene mitochondrial genomes from ^11^; the Okhotsk individuals were omitted from this analysis, due to their unique demographic history inferred from the PSMC. The bold line indicates the median value and the shaded area shows the 95% credibility intervals.

Investigations into the more recent maternal demographic history using mitochondrial Bayesian skyline plots indicate an increase in female N_e_ between 20-15 kya, followed by relative stability until ∼2.5 kya, when it begins to decrease until present (Fig 3B). This decrease is much sharper when including OKH individuals in the skyline analysis (Supplementary Fig. S10), but may be due to population structure in the dataset ^27^.

### Habitat connectivity

We calculated distances between bowhead whale suitable habitat patches within all four stocks throughout the Holocene at a fine resolution (see Methods), based on spatial analyses of projections from a bowhead whale ecological niche model ^13^ and satellite telemetry data ^12,26,28^ that indicated maximum bowhead movement limits.

Travel distances required to maintain connectivity between the BCB ↔ OKH populations regularly exceeded 4,500 km for the past 11,700 years (Fig. 4B). Given that these distances far exceed the upper observed seasonal movement distances for bowhead whales (∼3,500 km^12,29,30^), our habitat connectivity findings indicate that the Sea of Okhotsk bowhead whale population has been isolated throughout the Holocene.

**Figure 4:**
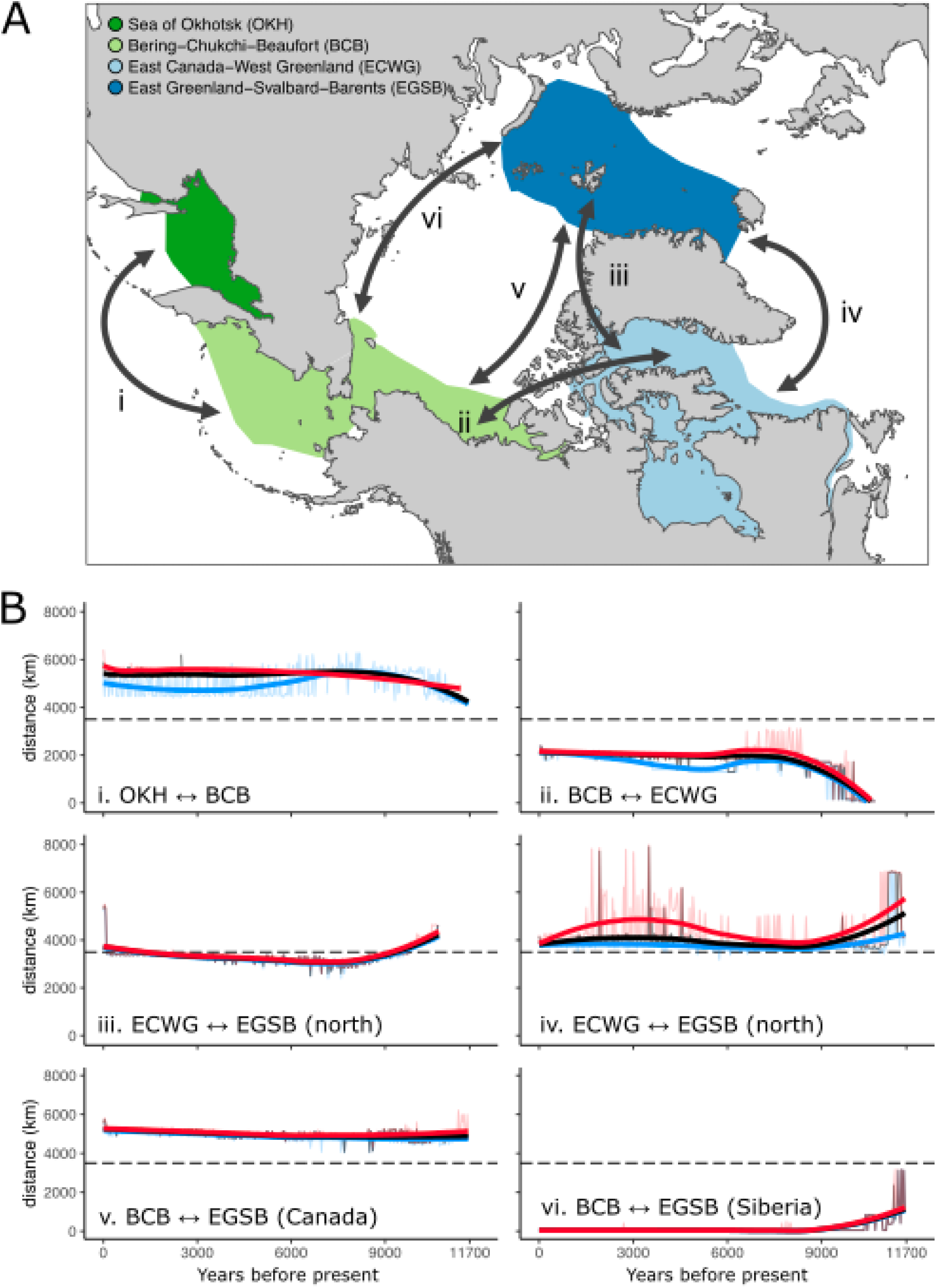
**Connectivity of bowhead whale populations across the Holocene**. **A)** Map of the four bowhead whale stocks. Arrows reflect potential dispersal pathways. **B)** Minimum inter-stock travel distances during the Holocene. Dashed black lines represent the 3,500 km upper bowhead whale movement estimate. Colors with associated 75% loess curves show different thresholds that exclude marginal bowhead whale habitat (blue = excluding lowest 10%; black = excluding lowest 12.5%; red = excluding lowest 15%).

In contrast, the closest core summer habitat patches between the BCB ↔ ECWG populations were rarely greater than ∼2,000 km once the Lancaster Sound in the Canadian Arctic Archipelago opened ∼10.7 kya ^31^ (Fig. 4). Thus, it can be expected that movement between these two populations was relatively frequent.

From ∼9 kya, when the Nares Strait that connects the Atlantic (northern Baffin Bay) and the Arctic (Lincoln Sea) Oceans opened ^32,33^, the nearest distance between the ECWG ↔ EGSB populations shortened to 3,000-4,000 km in length. This distance approximated the upper 3,500 km limit of bowhead whale movement, starting in the Holocene Thermal Maximum (10-5.5 kya ^34^) (Fig. 4). Distances between ECWG ↔ EGSB populations via a southern Greenland route mostly exceeded 3,500 km, only dipping below 3,500 km around ∼11 kya and again ∼5 kya. Distances between BCB ↔ EGSB habitat patches via Siberia were mostly <1,000 km throughout the Holocene. In contrast, BCB ↔ EGSB travel distances north of the Canadian Arctic Archipelago exceeded the 3,500km dispersal limit of bowhead whales.

Sensitivity analyses show that our estimates of inter-population connectivity are insensitive to thresholds used in the ecological niche modeling. We show that the exclusion of the most marginal habitats did not markedly increase travel distances between stocks (Fig. 4B). Furthermore, analysing spatial habitat suitability patterns through time showed that BCB ↔ ECWG and BCB ↔ EGSB travel distances were not sensitive to the exact location of population demarcation edges (SI Videos 1-3).

## Discussion

Understanding how past environmental and anthropogenic factors shaped contemporary genetic patterns is essential for assessing the vulnerability of species in the face of ongoing climate change ^35^. By integrating range-wide genomics with ecological modelling, we revealed how environmental change and habitat connectivity during the Holocene has shaped the genomic population structure and diversity of bowhead whales today. We identified three genetically differentiated bowhead whale populations, each showing little-to-no signs of recent inbreeding and high heterozygosity values relative to other mammalian species.

### Holocene Habitat Connectivity

Our nuclear genomic data indicate three genetically distinct populations, in contrast to the four management stocks currently recognised by the International Whaling Commission ^14^. We find the Sea of Okhotsk (OKH) bowhead whales are genetically distinct from all other stocks on a nuclear genomic level (Fig 1). Based on our demographic analysis, the OKH population began a unique trajectory, diverging from the others ∼200 kya (Fig. 3A). The relative isolation of the OKH bowhead whales is consistent with required long travel distances to the Bering-Chukchi-Beaufort (BCB) population (and all other populations via BCB), exceeding bowhead whale movement capacity of ∼3,500 km, at least for the past 11,700 years (Fig. 4). Thus, our findings indicate long-term geographic isolation has been a major driver of the pronounced genetic distinctiveness of the OKH population. However, the absence of pronounced phylogeographic structuring among the mitochondrial genomes suggests some level of connectivity via gene flow with the other populations (Fig. 1B and Supplementary Table S2).

Our data show that individuals from BCB and East Canada-West Greenland (ECWG) stocks are genetically indistinguishable (Fig. 1, Supplementary Figs. 5-8). Genomic similarity between these stocks is consistent with high habitat connectivity and short (≤2,000 km) inter-population travel distances throughout the Holocene (Fig. 4). It is also supported by contemporary research using satellite telemetry, which has demonstrated reciprocal movement between BCB and ECWG ^17^. Although this exchange was attributed to the recent and dramatic loss of sea ice in the Canadian Arctic, our results suggest connectivity between these regions may actually be a longer-term phenomenon, occurring at least since the opening of Lancaster Sound in the Canadian Arctic Archipelago ∼10.7 kya ^31^.

Our finding of individuals from East Greenland-Svalbard-Barents Sea (EGSB) forming a distinct cluster in the PCA (Fig. 1D), is supported by differences in stable isotope values between stocks, which indicate individuals predominantly foraged locally throughout the Holocene ^11^. However, ECWG and EGSB individuals have near-identical, deep-time demographic trajectories (Fig. 3A), suggesting they comprise a single population, or were distinct yet underwent similar demographic trajectories until ∼30 kya years ago, which is the approximate temporal limitation of the PSMC method ^36^. Time-series palaeogenomic data from Holocene bowhead whale fossils identified ECWG and EGSB to be panmictic throughout the past 12,000 years, possibly through low levels of migration (<5 individuals per generation) ^11^. The time-series data also showed that differentiation of the two stocks is a recent phenomenon, driven by commercial whaling, rather than climate and environmental factors ^11^. However, our habitat connectivity analysis indicates the distance between ECWG and EGSB was approximately at our estimated upper limit of bowhead whale dispersal (3,500 km) throughout the Holocene (Fig 4). Thus, we would expect that any dispersal between the populations would have been long-distance and at low frequency, supporting previous inferences ^11^.

Our habitat connectivity analyses identified that direct dispersal between the BCB ↔ EGSB populations via Siberian waters during the Holocene was plausible (Fig. 4). However, genetic evidence for a relatively strong connection between these stocks was not apparent (Fig 1D, Supplementary Fig S9). Although the coasts along Siberia have contained suitable habitat for bowhead whales throughout the Holocene (SI Videos 1-3), travel distance between the current designated stock boundaries would be ∼3,500 km one-way. With the exception of a single Late Holocene bowhead whale fossil from the East Siberian Sea ^37^, no bowhead whale fossils have (to our knowledge) been recovered between Franz Josef Archipelago and the Bering Strait ^38–41^. However, Arctic marine mammal fossils are rare in general from the Russian Arctic ^42^. Conservely, although contemporary aerial surveys north of Siberia are limited ^37^, bowhead whales have been detected in the region ^43^, supporting the capacity past connectivity between the BCB and EGSB populations, as suggested by our habitat analysis.

Our analyses could not identify whether north or south of Greenland has been a more likely dispersal path for bowhead whales between ECWG and EGSB in the past. While bowhead whale fossils dating to the Middle and Late Holocene have been recovered from the southernmost part of the Nares Strait and northeast of the Lincoln Sea further north, they are otherwise rare in northern Greenland ^42^. This is despite fossils of other Arctic marine mammals (narwhal, polar bear, ringed seal, walrus) having been found in the Nares Strait and Lincoln Sea, meaning that lack of preservation is not the cause for absence of recovered bowhead whale fossils in this area (Supplementary Fig S11). In contrast, in southeast Greenland, there is a scarcity of fossils from all Arctic marine mammal species ^42^. This may be due to acidity of substrates in the region, which likely limits preservation of any organic material ^11^.

Population genetic data are increasingly used to inform Inuit subsistence offtake quotas and stock status assessments ^14^, and genome-wide datasets offer a more powerful and nuanced conservation baseline for organisations such as the International Whaling Commission. The lack of genetic differentiation and historic movement barriers between the BCB and ECWG stock boundaries lends support towards treating them as a single management unit.

### Genomic health

Bowhead whales were nearly driven to extinction during four centuries of commercial whaling ^6^. Despite a long history of over exploitation, the BCB and ECWG management stocks are now relatively abundant, with present-day population sizes estimated at ∼16,000 ^44^ and ∼6,000 individuals ^44^, respectively. Less is known about the census size of stocks in the EGSB region and the Sea of Okhotsk, but they are each estimated to number in the low hundreds ^44–46^. Therefore, patterns of genetic diversity across bowhead whale populations may also partly reflect shorter-term anthropogenic impacts, as well as longer-term influences of environmental change and connectivity.

We identified three genetically well-differentiated bowhead whale populations, and each displayed high heterozygosity values relative to other mammalian species (Fig. 2B). The investigated individuals also had little-to-no signs of recent inbreeding (runs of homozygosity >1 Mb). Mitochondrial demographic reconstructions did not show a recent rapid decrease in female effective population size that could be associated with commercial whaling (Fig. 3B). While the large population declines associated with commercial whaling would be expected to lead to reduced genetic diversity, increased inbreeding, and decreased nuclear effective population size, this pattern is not clearly observed. This absence may reflect time lags between demographic and genetic changes^47^, as has been shown for ECWG and EGSB ^11^.

The higher genome-wide heterozygosity levels in BCB/ECWG may reflect a larger historical population size and higher long-term diversity (Fig. 2A). The Sea of Okhotsk had the lowest diversity estimates (Fig. 2A,C), which likely reflects the long-term isolation and putative long-term low population size of the population.

Despite having undergone one of the most dramatic reductions in population abundance due to historical whaling (>90% decline) ^11^, the EGSB population retains high genetic diversity relative to the other bowhead populations (Fig. 2A,C), perhaps due to high pre-whaling abundances and/or diversity. EGSB is estimated to have numbered in excess of 52,500 individuals prior to commercial whaling ^48^, and palaeogenomes have shown pre-whaling genetic diversity was slightly higher (∼2%) than today ^11^. However, their apparent high present-day diversity hides underlying genetic vulnerabilities. Our results show that EGSB bowhead whales harbour the highest masked genetic load, with a substantial proportion of deleterious variants present in the heterozygous state (Supplementary table S4). Bowhead whales in the EGSB stock may become more vulnerable as the genomic signatures of their past demographic collapse become evident. For example, if the population decline leads to increased inbreeding in the future, many of the currently masked deleterious mutations may become homozygous, increasing the realised genetic load and posing a risk to population viability and fitness.

The OKH population, shaped by long-term isolation, exhibits both reduced genetic diversity and elevated realised load (Fig. 2 and Supplementary table S4)—consistent with the prolonged effects of genetic drift and limited gene flow. The effects of climate change on the population will be all the more severe, given that the Sea of Okhotsk is already ice-free during the summer ^13^. Bowhead whales in this region never appear to leave the Sea of Okhotsk ^49^. This is because traveling around southern Kamchatka and into the Bering Sea may also increase their exposure to attacks by killer whales (*Orca orcinus*) ^50^. Future predictions of suitable bowhead whale habitat based on the ecological models used here indicate the complete loss of suitable habitat in the Sea of Okhotsk by 2060 CE ^13^. In combination with our findings of lower genetic diversity and higher inbreeding levels, the OKH population is thus the most vulnerable to extirpation in the near future.

Our various lines of evidence illustrate that while the natural environment may have structured genetic variation in bowhead whales across longer time scales, anthropogenic impacts of the past 500 years may have shifted populations from historically stable evolutionary trajectories toward increased genetic risk—underscoring the need for timely, population-specific conservation strategies.

### Future outlook

Looking to the future, continued climatic change and other human-driven threats may also reshape population connectivity. Bowhead whales are now spending winters further north – such as in the southern Chukchi Sea – in response to sea ice decline ^51^. They are more ice-associated during the winter than in the summer ^26,52^, possibly as this reduces migration distance to their primary summering grounds ^53^. Our connectivity analysis focused on summer connectivity because it is the main foraging period of bowhead whales ^54^. Continued loss of Arctic sea ice could facilitate new dispersal routes between currently separated populations—for instance, increased movement along the Siberian coast, around northern Greenland, or across the polar basin. While such changes may enable gene flow and buffer small populations against isolation, they may also introduce new threats, including: increased disease risks; increased exposure to shipping traffic ^55^; increased summer overlap with predatory killer whales ^56^; competition with sub-Arctic cetacean species moving their distributions further north ^57^.

As Arctic sea ice continues to decline and human activity intensifies across the region, the future dispersal potential and adaptive resilience of bowhead whale populations will depend not only on their environmental responses (and those of their prey), but also on their demographic histories and genomic health. In this context, conservation planning must account for both historical and future connectivity, the complex interplay between demography and genetic load, and the risks posed by climate-driven shifts in Arctic marine ecosystems. By integrating range-wide population genomics with spatially-explicit habitat connectivity modelling, this study not only improves our understanding of bowhead whale population dynamics, but also provides a scalable framework for predicting the evolutionary and conservation trajectories of other Arctic marine species facing the dual pressures of historical exploitation and accelerating environmental change.

## Methods

### Genetic data

#### Samples

We used samples from contemporary bowhead whale individuals collected across the distribution range of the species. Our samples represent all four recognised management stocks (Fig. 1A, Supplementary table S1): East Greenland-Svalbard-Barents Sea (EGSB); Eastern Canada-West Greenland (ECWG); Bering-Chukchi-Beaufort Seas (BCB); and the Sea of Okhotsk (OKH) ^16^.

We analysed nuclear genomes from 43 individuals, 24 of which are new to this study. We removed two individuals identified as close relatives (see details below) and two others with very low coverage (<0.005x), basing our population analyses on the remaining 39 nuclear genomes. We also analysed 48 mitochondrial genomes (mitogenomes), of which 43 overlapped with the nuclear samples. The novel data from each stock were: ECWG (*n* = 7); BCB (*n* = 7); and OKH (*n* = 10) (Supplementary table S1). OKH samples were imported with CITES Permit №21RU000001 (issued 11.01.2021), while all other samples were imported under CITES exemption DK014 to the Globe Institute, University of Copenhagen.

We sourced published bowhead whale genetic data from various publications ^11,23,24,58,59^. Despite the availability of sequencing data for additional EGSB individuals B, C, D, E, and F, we excluded them because they might be duplicate samples of the individual ‘A’ ^23,58^.

Based on our initial analyses of nuclear population structure, which suggested the BCB and ECWG individuals represented a single population, we used high coverage (>20x) representative individuals from each genetic population for individual-based analyses of diversity, demography, and genetic load. Two individuals with newly generated data were from Igloolik, Central Canada (CGG_1_024320), and the Sea of Okhotsk (CGG_1_024474), which we used to represent the BCB/ECWG and OKH populations respectively. We used published data from the individual 17-19, sampled in the Greenland Sea between Greenland and Svalbard, to represent the EGSB population ^23^.

#### Data generation

For the 24 individuals new to this study, we extracted DNA from tissue samples using a DNeasy blood and tissue kit following the manufacturer’s protocol. We fragmented the extracted DNA from the new ECWG and BCB samples to ∼450 bp using a Covaris M220 ultrasonicator. We built Illumina sequencing libraries from the sonicated DNA using the BEST protocol ^60^, using Illumina adapter mix concentration of 20uM and 15 cycles during the indexing PCR step. We cleaned the indexed libraries using a SPRI bead DNA purification method. For the OKH samples, the extracted DNA was shipped to Novogene (UK), fragmented, and built into Illumina sequencing libraries. All samples were sequenced on an Illumina Novaseq 6000 at Novogene using 150 bp PE reads. DNA previously extracted for individual CGG_1_24320^11^ was sent to StarSEQ, Germany, to have additional Illumina sequencing libraries constructed and sequencing performed to ∼25x coverage using an Illumina Nextseq 2000.

#### Data processing

We trimmed adapter sequences and removed reads shorter than 30 bp from the raw reads using skewer v.0.2.2 ^61^. We merged overlapping paired-end reads using FLASHv1.2v11 ^62^, using the default parameters. We mapped the merged and unmerged reads to the bowhead whale reference genome ^24^ including the mitochondrial genome (Genbank accession: KY026773.1) using Burrows-wheeler-aligner (BWA) v0.7.15 ^63^, the mem algorithm, and default parameters. We parsed the output and removed duplicates and reads of mapping quality <30 using SAMtools v.1.6 ^64^. Mapping to the reference bowhead whale nuclear genome resulted in genome-wide coverages between 0.001x and 26.55x (median = 3.27x, mean=6.44x, Supplementary Table S1). Two individuals, CGG_1_024323 and CGG_1_024325, both from the BCB stock, had very low nuclear genome coverage (<0.005x) and were removed from downstream nuclear genomic analyses. Mapping to the reference bowhead whale mitochondrial genome resulted in mitochondria-wide coverages between 97.3x and 7920.9x (median=1631.3x, mean=2859.0x).

#### Sex scaffold identification

We identified putative sex chromosome scaffolds in the bowhead whale reference genome by aligning it to the Cow X (Genbank accession: CM008168.2) and Human Y (Genbank accession: NC_000024.10) chromosomes. We performed the alignments using satsuma synteny v2.1 ^65^ with default parameters.

#### Relatedness

We assessed the nuclear genomic relatedness of all 41 individuals with sufficient coverage (>0.2x) using NGSrelate v2 ^66^. As input, we calculated genotype likelihoods using ANGSD v0.921 ^67^. We obtained genotype likelihoods using the GATK algorithm (-GL 2), specified the output as a binary beagle file (-doGlf 3), and applied the following filters: only include reads with a mapping and base quality greater than 20 (-minmapQ 20 -minQ 20); only include reads that map to one location uniquely (-uniqueonly 1); a minimum minor allele frequency of 0.05 or greater (-minmaf 0.05); only call a SNP if the p-value is less than 1e^-6^ (- SNP_pval 1e-6); infer major and minor alleles from genotype likelihoods (-doMajorMinor 1); skip triallelic sites (-skipTriallelic 1); remove sex scaffolds and scaffolds shorter than 100 kb (-rf); and call allele frequencies based on a fixed major and an unknown minor allele (-doMaf 2). We determined a relatedness coefficient (RAB) >0.25, equivalent to at least first cousins, as closely related and we removed whichever individual had the lowest coverage from downstream nuclear genomic analyses. This led to the exclusion of two individuals; CGG_1_024313 from ECWG and CGG_1_024482 from OKH. We also obtained inbreeding coefficients (F) for each individual from the NGSrelate output.

Individuals DL7642 and CGG_1_024313 had a RAB relatedness score of 0.44, and individuals CGG_1_024478 and CGG_1_024482 had a RAB relatedness score of 0.26. We therefore removed the individuals with the lowest coverage (CGG_1_024313 and CGG_1_024482; 1.3x and 3.3x) from each pair for further analyses.

#### Population structure

We used both pseudohaploid consensus base calls and genotype likelihoods for the PCA. We generated the pseudohaploid identity-by-state base calls and genotype likelihoods in ANGSD (Korneliussen et al., 2014) using the following filters and parameters: -uniqueOnly 1, -GL 2, -remove_bads 1, -minMapQ 20, -minQ 20, -SNP_pval 1e-6 -skipTriallelic 1, -doGlf 2, - domajorminor 1, -minmaf 0.05, minimum number of individuals required to consider the site in the analysis as 20 (-minind 20), compute pseudohaploid consensus base calls (-doIBS 2), only consider a site if the minor allele is present in at least two individuals (-minminor 2), and output a covariance matrix from the identity-by-state base calls (-doCov). Additionally, we constructed a covariance matrix from genotype likelihoods using PCAngsd v0.98 (Meisner & Albrechtsen, 2018). We determined the significance of PCA axes using a Tracy-Wildon test on the eigenvalues in R^68,69^ based on a significance value of p < 0.01.

To build a neighbour joining tree, we generated a pairwise distance matrix from ANGSD when performing the pseudohaploid base calls (setting the parameter -makematrix 1). We converted the distance matrix into a neighbour-joining phylogenetic tree using fastME v2.1.6.1 ^70^ and default parameters.

We calculated admixture proportions using the same genotype likelihoods calculated for the PCA with NGSadmix (Skotte et al., 2013). We ran NGSadmix specifying K = 2 and K = 3. To evaluate the reliability of the NGSadmix results, we independently ran each K up to 50 times. We considered K values reliable if we retrieved consistent log-likelihoods from at least two independent runs.

For the fixation index, we ran ANGSD with the same filtering parameters, but with the addition of calling consensus pseudohaploid bases (-dohaplocall 2). We calculated *F*_ST_ from the pseudohaploid base call file in 100 kb non-overlapping sliding windows, with a minimum requirement of 100 sites per window using the available popgenWindows.py (https://github.com/simonhmartin/genomics_general). To investigate the impact of differences in coverage between our sampleset, we repeated the analyses but with the 12 EGSB individuals downsampled to ∼2x using SAMtools.

We used D-statistics (ABBA BABA) to investigate gene flow between non-sister populations^71^. We calculated D-statistics for all individuals using a random base call approach in ANGSD (-doabbababa 1), specifying only autosomal scaffolds >100 kb with the following parameters: -minmapQ 20, -minQ 20, -blocksize 1000000, -uniqueonly 1. To infer the ancestral sequence (-anc) we used a right whale (*Eubalaena glacialis*) individual (BioSample: SAMN32746534), which we mapped to the bowhead whale reference genome using the same approach as for bowhead whales. We summarised the results using a block jackknife approach with the R script available in the ANGSD toolsuite (ANGSD_jackknife.R). We subsequently extracted only comparisons which fit the known population structure of our samples, i.e. ((ECWG, BCB), EGSB), ((BCB, ECWG), OKH), ((EGSB, BCB), OKH), and ((EGSB, ECWG), OKH). This allowed us to investigate whether there was relatively more gene flow between OKH and any of the remaining management stocks as well as between EGSB and either ECWG or BCB. We considered all Z-scores > |3| to be significant.

#### Genetic diversity

We calculated autosomal heterozygosity in one high coverage (>20x) individual per population using ANGSD: 17-19 (20.94x, EGSB, CGG_1_024320 (26.55x, ECWG/BCB), CGG_1_024474 (20.30x, OKH). To ensure comparability with previous studies, we followed the filtering parameters of Cerca et al. ^23^. These filters included: use the SAMtools algorithm to calculate genotype likelihoods (-GL 1); the minimum read depth for a site to be 5 (- setMinDepthInd 5); -minmapq 25, -minq 25; -uniqueonly 1; -remove_bads 1; calculate sample allele frequencies (-doSaf 1); use the reference genome as the ancestral sequence (- anc); only consider autosomal scaffolds >100 kb (-rf); adjust quality scores around indels (- baq 1); and fold the site frequency spectrum (-fold 1). We called the folded SFS from the allele counts using realSFS, part of the ANGSD toolsuite, in 100 kb non-overlapping sliding windows. We generated the 100 kb sliding windows bed file using bedtools ^72^. We obtained mean nuclear genome-wide heterozygosity values from 25 other mammalian species for comparison from Cerca et al. ^23^.

We calculated population levels of nucleotide diversity from the pseudohaploid base call file generated for the *F*_ST_ analysis in 100 kb non-overlapping sliding windows, with a minimum requirement of 100 sites per window using the available popgenWindows.py (https://github.com/simonhmartin/genomics_general). Based on the population structure results, we pooled individuals into three populations, EGSB, OKH, and ECWG/BCB. To correct for differences in coverage, we repeated this analysis with the 12 EGSB individuals downsampled to 2x, and calculated mean differences between the original and downsampled versions. We then used that to correct for the nucleotide diversity in remaining stocks due to lower-coverage individuals.

To investigate inbreeding we used ROHan ^73^ to calculate runs of homozygosity (ROH). We ran ROHan using default parameters, but specified a window as being a ROH if it had an average heterozygosity of < 5x10^-5^, as opposed to 1x10^-5^, and limited the analysis to only autosomal scaffolds >1 Mb.

#### Genetic load

We calculated the genetic load of the three high-coverage individuals using SnpEff v5.1 ^74^. As input, we created a VCF file containing only variant positions using BCFtools v1.15^75^ for the autosomes >100 kb and with the minimum mapping and base quality filters set to 20 (-q 20 -Q 20). This VCF also included the *Eubalaena glacialis* individual mapped to the bowhead whale reference genome, which we also used in the D-statistics analysis.

We built a SnpEff database using the basic config file, but with the addition of the bowhead whale reference genome and its annotations using the build function. We filtered the annotated output VCF based on three groupings: loss of function (LoF), nonsynonymous mutation, and synonymous mutation. We designated SNPs as LoF if the site was annotated as “stop_gained”, “frameshift_variant”, ”splice_acceptor_variant”, or “splice_donor_variant”. We designated SNPs as nonsynonymous if a site was annotated as “missense_variant”, and synonymous if a site was annotated as either “synonymous_variant”, ”start_retained” or “stop_retained_variant”.

We further filtered for sites where the alternative allele was designated as a LoF or nonsynonymous variant and *Eubalaena glacialis* was homozygous for the reference allele. If a site was homozygous, we only considered this of interest if the bowhead individual had the derived allele - the *Eubalaena glacialis* allele was considered the ancestral allele. To normalise our values, we divided the number of LoF or nonsynonymous sites by the number of synonymous variants relative to the reference allele. We considered a heterozygous synonymous variant as a single change and a homozygous synonymous variant as two changes.

#### Demographic history

We ran demographic analyses on diploid consensus genomes from the three high-coverage individuals using PSMC ^36^. We called diploid genome sequences using SAMtools and BCFtools, specifying a minimum quality score of 20, minimum coverage of 10, and specified only to autosomes >100 kb in length. We ran PSMC specifying standard atomic intervals (4+25*2+4+6), a maximum number of iterations of 25 (-N), maximum 2N0 coalescent time of 15 (-t), and initial theta/rho ratio of 5 (-r), and performed 50 bootstrap replicates. We checked PSMC results for overfitting so that after 20 iterations, at least 10 recombinations were inferred to have occurred in the intervals each parameter spans. We plotted the output using a bowhead whale mutation rate of 1.2x10^-8^ per generation ^76^, assuming a generation time of 35 years ^18^.

#### Mitochondrial genomes

We built consensus sequence mitochondrial genomes using a majority rules approach (- doFasta 2) in ANGSD only considering bases with quality scores >30 (-minq 30), reads with a mapping quality scores >30 (-minmapq 30), and sites with at least 10x coverage (- minInddepth 10). We aligned the mitochondrial genomes using Mafft v7.310 ^77^ and default parameters.

We constructed an unrooted haplotype network for the 48 complete mitochondrial genomes of the bowhead whale using the Median-joining network ^78^ as implemented in PopART ^79^. We performed stock diversity estimates with DnaSP v.6.12.03 ^80^. Diversity statistics included the number of haplotypes (*h*), the number of segregating sites (*S*), the haplotype (H_D_) and nucleotide (π) diversity ^81^ and Tajima’s D ^82^. Gaps and missing data were considered and excluded only in pairwise comparisons.

We interred the demographic history of bowhead whale mitochondrial genomes by employing the Bayesian skyline plot method ^83^ implemented in BEAST v.2.6.1 ^84^ using all mitochondrial genomes generated in this study and 107 previously published radiocarbon dated ancient individuals from the Canadian Arctic Archipelago and Svalbard regions ^11^. We repeated this analysis twice. One run included all individuals (*n* = 155). Then we ran the analysis without the 11 Okhotsk individuals to avoid the effects of population substructure, as the OKH stock was relatively distinct to the other populations ^27^. As input, we aligned the mitochondrial genomes using Mafft v7.392 ^77^ and extracted 38 regions, including protein-coding regions, rRNAs, tRNAs and control region, based on GenBank annotation coordinates. We combined sequences of these 38 regions into six subsets: (i) first codon position of the protein-coding regions; (ii) second codon position of the protein-coding regions; (iii) third codon position of the protein-coding regions; (iv) tRNAs; (v) rRNAs; and (vi) control region. We identified the best-fit partitioning scheme and substitution model for the six subsets using Partitionfinder v.2.1.1 ^85^. The best partitioning scheme and substitution models, based on the corrected Akaike Information Criterion, were used as input for Beast2. We analysed each partition using unlinked substitution models, but linked genealogy and molecular clocks. We assumed six groups of coalescent intervals and a strict molecular clock with a rate of one unit per substitution per site. We sampled from posterior distributions of parameters using MCMC methods. MCMC sampling consisted of 100 million burn-in steps followed by 1,000 million steps, sampled at every 100,000 steps. We assessed chain mixing and convergence to stationarity using Tracer v.1.7.1 ^86^, and required minimum effective sample sizes of 5,000 for all parameter estimates.

#### Habitat Connectivity

We leveraged summer habitat suitability generated from an ecological niche model (ENM) of bowhead whales at 35-year time steps (corresponding to the average generation time of the species ^87^) throughout the Holocene ^13^. This ENM was built using occurrence data from fossils, historical whaling records, as well as contemporary sightings and satellite tracking (see supporting information of ref ^13^). This ENM was projected at a 60 km x 60 km pan-Arctic projection, as this spatial scale allowed for accurate representation of the opening of known channels (e.g., Lancaster Sound, Nares Strait) due to sea-level rise and ice-sheet deglaciation (see next paragraph). Occurrence data was matched in time and space to modelled palaeoclimate reconstructions from a HadCM3B-M2.1 coupled atmosphere-ocean general circulation model ^88^. Cross-validation tests show that the ENM did well at predicting occurrence as a response to climatic and environmental conditions, and also identified known bowhead whale summer congregation hotspots ^13^.

To account for sea-level dynamics operating at resolutions finer than our ENM projections, we generated a high-resolution, temporally changing, land-sea mask for the Holocene. To do this, we statistically downscaled ICE-7G_NA deglaciation reconstructions ^31^ from a 1° x 1° resolution to a 5 km x 5 km resolution using GEBCO 2014 bedrock bathymetry data (https://www.gebco.net/) ^89^ via the delta-method (Supplementary Figure S1) ^90^. Reconstructions from the ICE-7G_NA model are available at 500-year time steps for the Holocene, which we bilinearly interpolated to the 35-year time steps used in the ENM. To identify and mask out potential dispersal corridors that would not have been traversable by bowhead whales due to being glaciated, we used fine-scale (rasterized to 5km x 5km) ice sheet margin (i.e., edge) reconstructions for: the Laurentide, Cordilleran, & Innuitian ^91^; Greenland and Iceland ^92^, Fennoscandian ^93^; and Eurasian (minus the Fennoscandian) ice sheets ^94^. We treated areas above sea level and areas covered by ice sheets as impenetrable. We then upscaled these fine resolution ice sheet-land-sea masks to the ENM resolution where a 60 km x 60 km grid cell was classified as land/ice sheets if its centroid was covered by the 5 km x 5 km mask (Supplementary Figure S1). This approach retained the finer resolution dispersal channels without enabling dispersal across impenetrable barriers like isthmuses (e.g., northern Kamchatka).

We calculated connectivity between the four bowhead whale stocks throughout the Holocene using the landscapemetrics and terra R packages ^95,96^. We did this by first identifying “core patches” of summer habitat (grid cells with all eight surrounding cells also being suitable; Supplementary Figure S2) and calculating pairwise travel distances between all core patches, accounting for temporally evolving land and ice sheet barriers. We used core habitat patches, rather than single cells, to better capture spatial population dynamics and to exclude potential edge effects (Supplementary Figure S3) ^97^. We identified minimum travel distances (inter-stock travel distance) between core patches of all four bowhead whale stocks (Supplementary Figure S3).

We calculated the following inter-stock travel distances: (i) OKH ↔ BCB; (ii) BCB ↔ ECWG; (iii) ECWG ↔ EGSB via Lancaster Sound/Nares Strait; (iv) ECWG ↔ EGSB via south of Greenland; (v) BCB ↔ EGSB via the northern edge of the Canadian Arctic Archipelago/Greenland; and (vi) BCB ↔ EGSB via the northern edge of Siberia. We included core patches outside contemporary management stock boundaries to account for potential dispersal corridors that existed during the Holocene and have since been lost due to climate change and effects of commercial whaling ^14^. For ECWG ↔ EGSB and OKH ↔ BCB travel distances, we selected the northern/southern edge of Greenland and the southern edge of Kamchatka, respectively, to demarcate if an area was closer to a given population, as marine species would have to pass through these areas to minimize travel distance between stocks. Thus, these land masses reflect natural biogeographic barriers to marine species. For BCB ↔ ECWG and BCB ↔ EGSB travel distances, given there were no obvious biogeographic barrier choke-points that minimized distance between these populations, we defined the halfway distance between contemporary management stock boundaries as a demarcation line for stock affinity.

We investigated whether more marginal (low-quality) habitat substantially shortened inter-population travel distances throughout the Holocene (Supplementary Figure S4). We did this by setting higher occurrence thresholds for occurrence in our ENM projections ^98^. We used the “10% occurrence” threshold as in ^13^ as well as 12.5% and 15% occurrence thresholds. Setting thresholds >15% resulted in the near absence of nearly all core patches (and suitable habitat in general) in the Sea of Okhotsk throughout the Holocene, which is unlikely, given the region’s whaling history and present-day persistence of bowhead whales.

We compared our estimates of inter-stock travel distances to observed bowhead whale movement distances using available tracking data for 83 individuals, of which two tracks are new to this study ^28^, totalling 53,955 geolocations ^12,26^. We removed position reports that were classified as poor ^99^ and calculated cumulative bowhead whale travel distance as a function of total track days. We estimated bowhead whale cumulative travel distance to be ∼7,000 km*yr^-1^, meaning that possible seasonal one-way travel distances are ∼3,500 km (Supplementary Figure S4). These distances are slightly inflated due to the inclusion of daily foraging patterns not related to directional movements. Calculating maximum travel distance (around land barriers) between known bowhead summering and wintering grounds revealed similar numbers, with distances of ∼2,500 km and ∼3,200 km in the ECWG and BCB stocks respectively ^12,29,30^. Therefore, 3,500 km represents a reasonable upper limit for bowhead seasonal movement distances. If the bowhead whale stocks were consistently further apart than this upper observed movement distance, dispersal between stocks would be far less likely, and would lead to genetically distinct populations forming over time due to isolation.

## Acknowledgements

Thanks to the hunters that have contributed samples. We thank Kristin Kaschner and Cristina Garilao for assistance with early suitable habitat models. Funded by Villum Fonden YIP+ grant no 37352 and DFF Sapere Aude grant no 9064-00025B to EDL. The Norwegian Research Council (through the ICE-whales project 244488/E10), the Greenland Institute of Natural Resources, the Environment in the North funding programme from the Department of Foreign Affairs (Norway) and the Norwegian Polar Institute supported the EGSB collections and some laboratory work.

## Data availability

Raw sequencing reads for all newly generated genomic data will be available on Genbank by the time of acceptance.

