## Supplementary information for "Tracing the drivers of range-wide bowhead whale genomic structure and diversity"

### Supplementary Figures

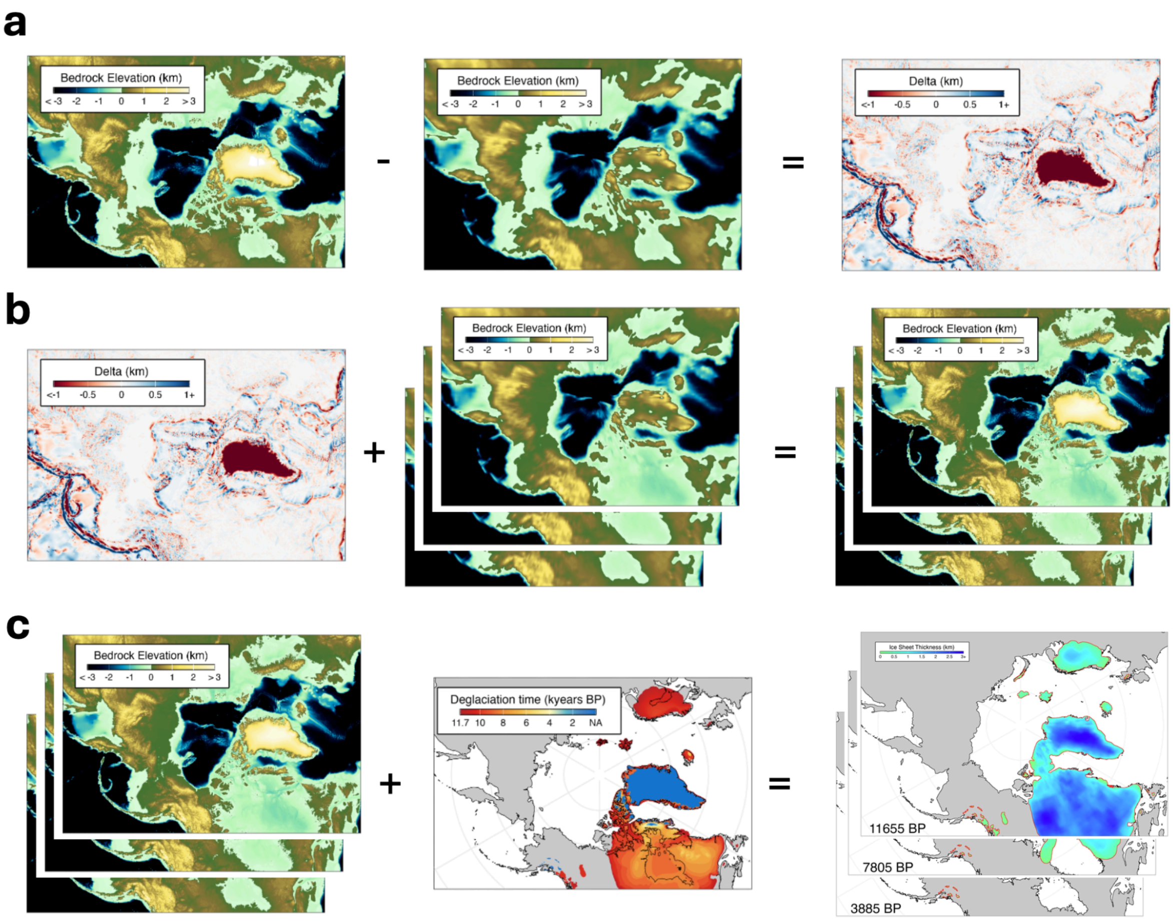


**Supplementary Figure S1 – Generating a fine-scale spatiotemporally varying land-sea mask via statistical downscaling**. **a.)** The contemporary bedrock elevation data of the ICE-7G_NA deglaciation model (which has been bilinearly interpolated to 5 km x 5 km resolution from its native 1° x 1° resolution) is subtracted from the fine-scale (5 km x 5 km) contemporary bedrock elevation GEBCO data to produce a delta layer containing fine-scale anomalies between the two layers (i.e., an additive delta method). **b.)** This delta layer is then added to the ICE-7G_NA bedrock layers (which have also been bilinearly interpolated to 5 km x 5 km from their native 1° x 1° resolution) for all the Holocene time bins to produce fine-scale bedrock elevation reconstructions. This assumes that the geometry of fine-scale structures (e.g., sub-1° features such as subterranean canyons, small bays/fjords, etc.) are preserved over the course of the Holocene. **c.)** Areas above sea level and areas covered by ice sheets are then masked out to produce fine-scale reconstructions of the land-sea mask. These are then upscaled to the 60 km x 60 km resolution used in analyses.

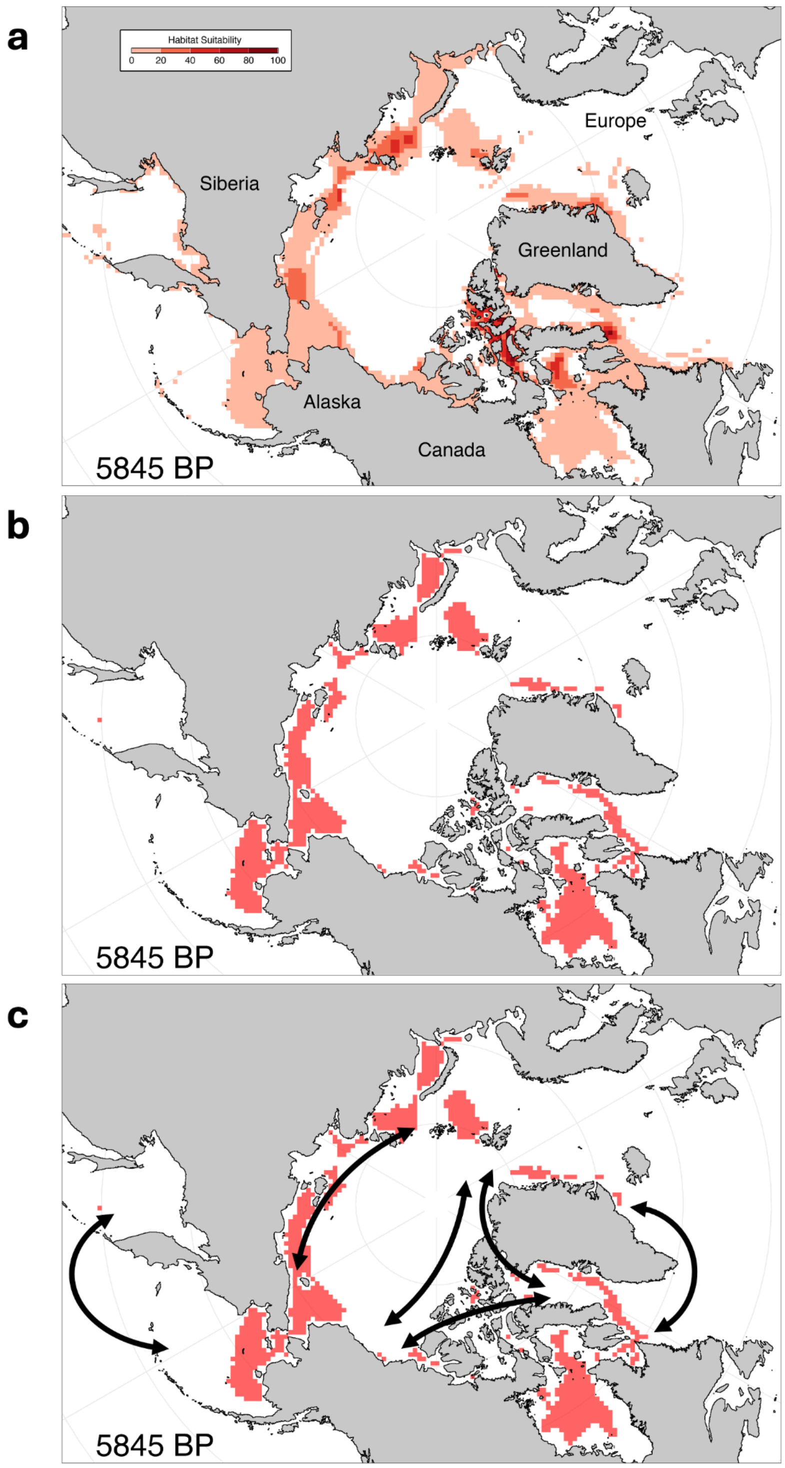


**Supplementary Figure S2 – Identifying core habitat patches and quantifying inter-stock travel distances**. **a**.) Thresholded habitat suitability projections from the bowhead whale niche model. Areas on land or covered by ice sheets have been removed according to the spatio-temporally varying land-sea mask (e.g., 5845 years BP). **b**.) Core habitat patches (suitable grid cells surrounded on all sides by other suitable grid cells) are identified. **c**.) The minimum inter-stock travel distances are calculated accounting for the spatio-temporally varying land-sea mask. This process is then repeated for all other time bins and all other niche model-threshold values.

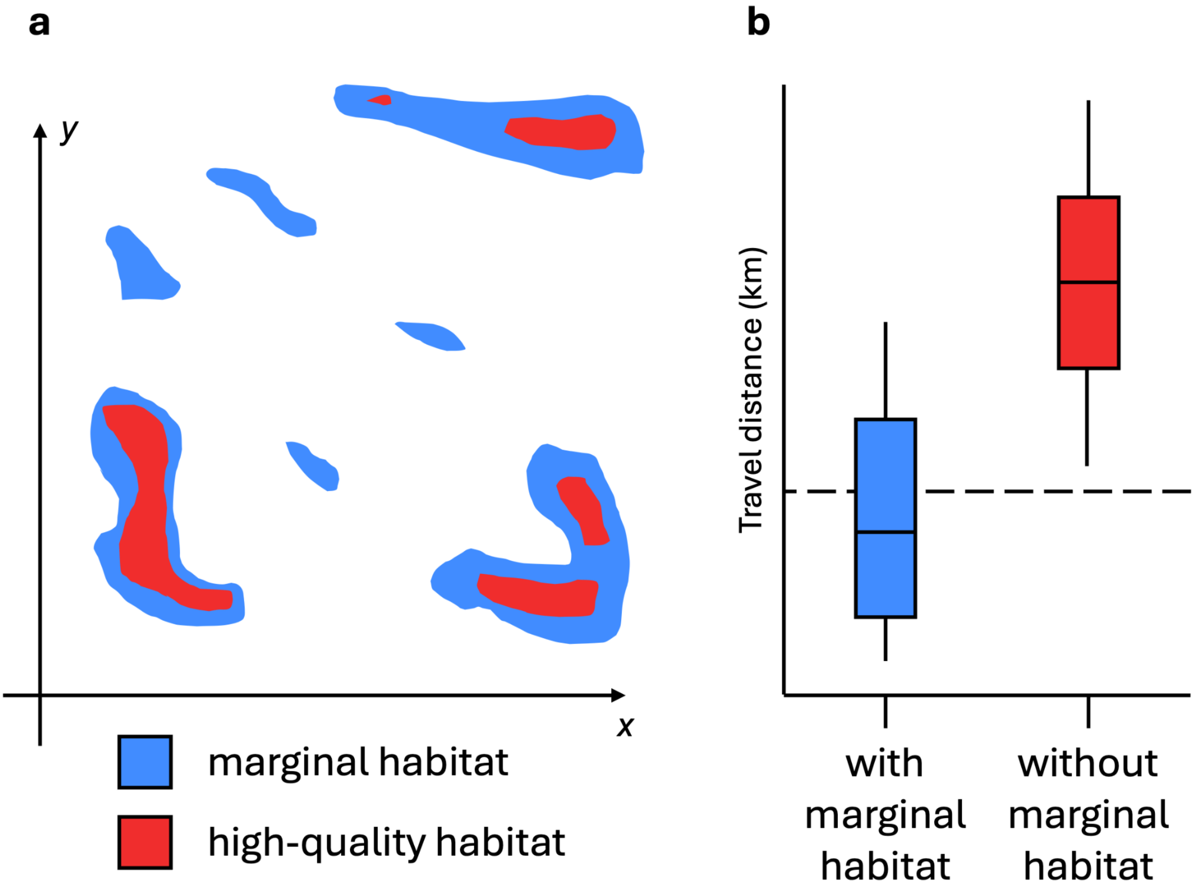


**Supplementary Figure S3 – Rationale for evaluating the role of marginal habitats in shortening inter-stock travel distances**. **a**.) Theoretical 2-dimensional representation of a sea-scape with marginal- (blue) and high quality-habitat core patches (red). **b.)** Boxplots of pairwise distances between habitat patches when including vs omitting marginal habitat. The dashed line represents a theoretical maximum bowhead whale travel distance. In this hypothetical example, marginal habitat patches may not sustain many individuals, but they may serve as valuable stopover locations, markedly lowering travel distances between habitat patches, ultimately maintaining meta-population connectivity over long time periods.

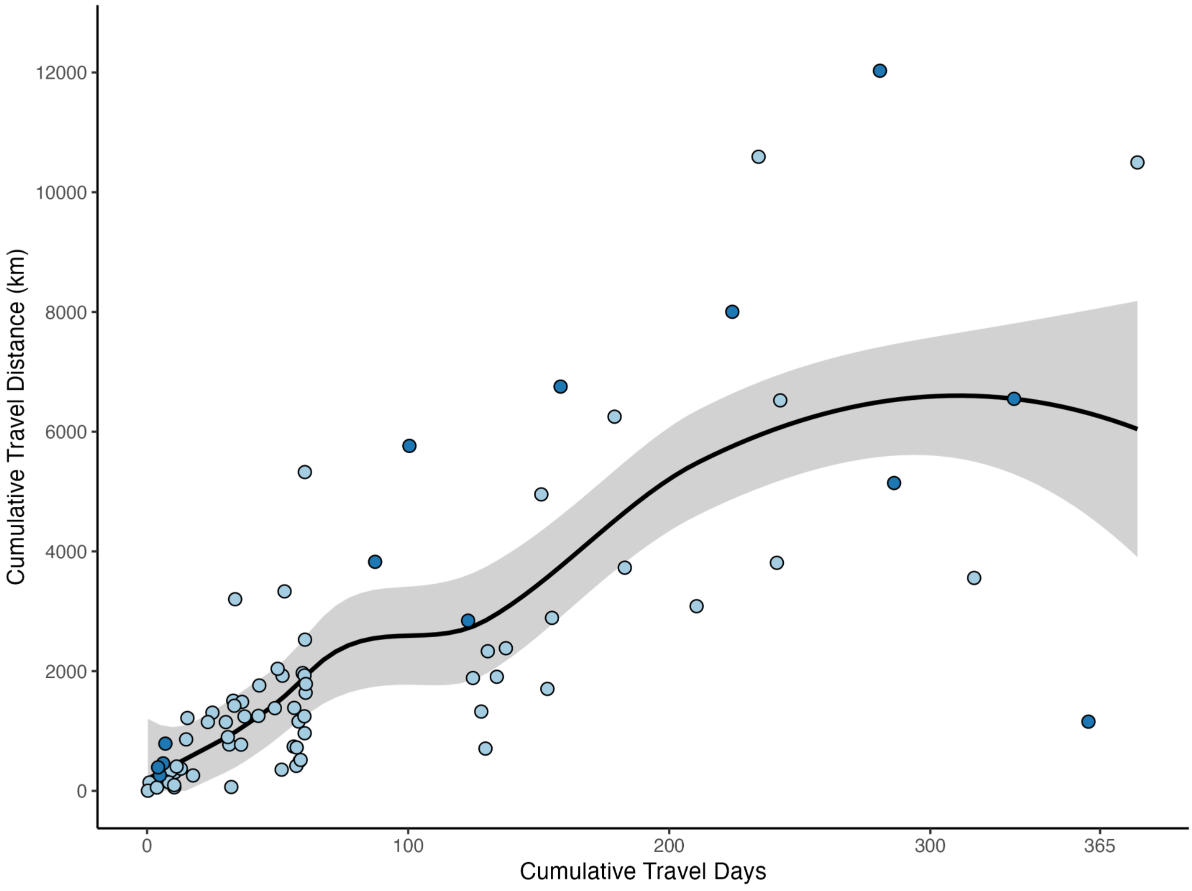


**Supplementary Figure S4 – Migration distances for bowhead whales**. Cumulative travel distances as a function of the number of days a satellite tag was attached to an individual whale. Only one-third of all tags (*n* = 81) were attached longer than 60 days. The curve represents a loess regression of all tracks. Colors represent different bowhead whale populations: East Greenland-Svalbard-Barents = dark blue, *n* = 13; East Canada-West Greenland = light blue, *n* = 68. No raw tracking data for bowhead whales in the Bering-Chukchi-Beaufort was publicly available, and no satellite tagging data exists for bowhead whales from the Sea of Okhotsk.

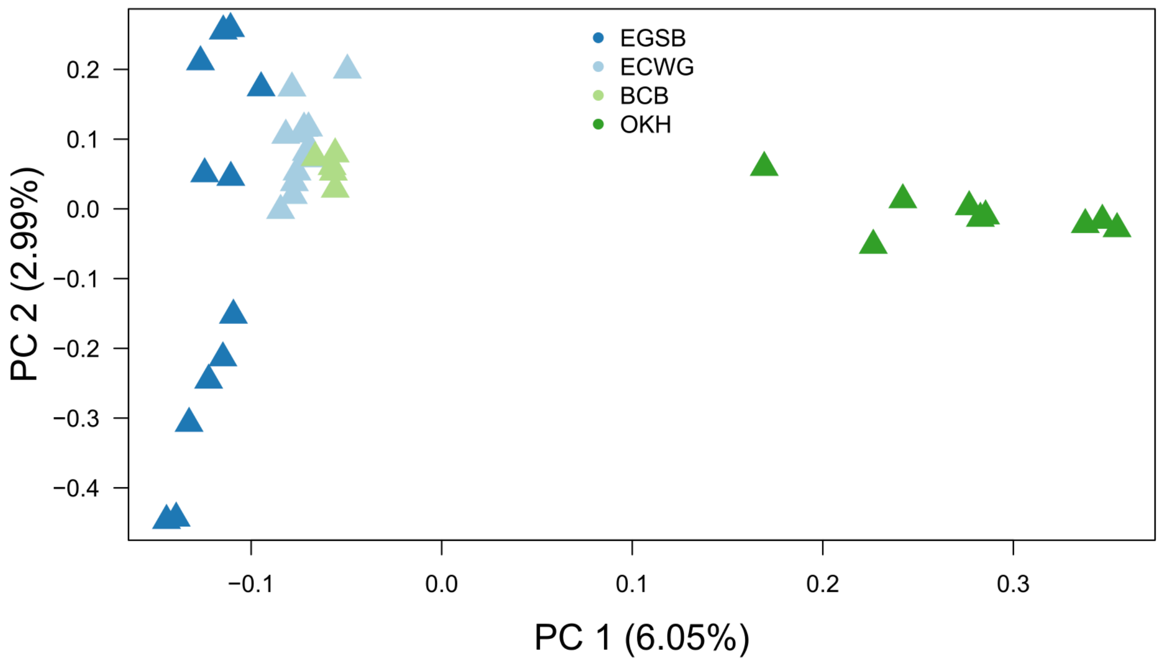


**Supplementary Figure S5:** Nuclear genomic PCA created from using genotype likelihoods. Percentages of variation explained by the PC 1 and PC 2 are indicated in parentheses.

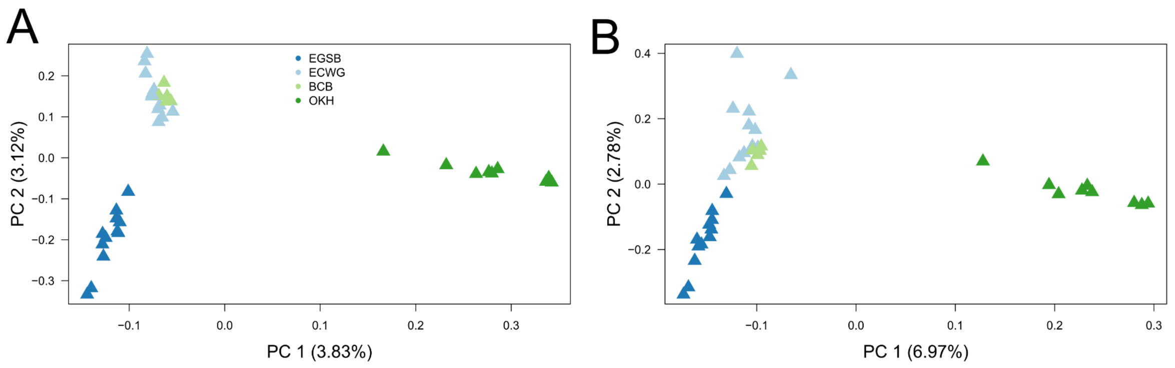


**Supplementary Figure S6:** Nuclear genomic PCA created from using **A)** pseudohaploid basecalls or **B)** genotype likelihoods and having the EGSB individuals downsampled to 2x to be more comparable to the other individuals. Percentages of variation explained by the PC 1 and PC 2 are indicated in parentheses.

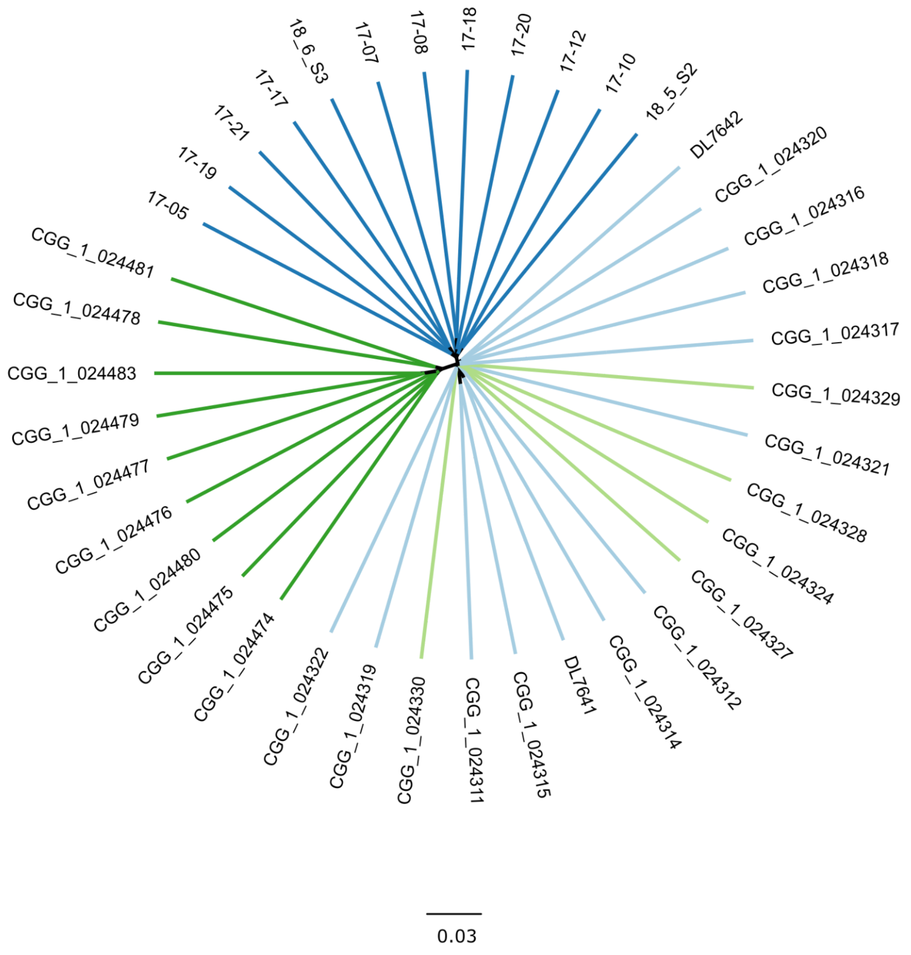


**Supplementary Figure S7:** Neighbour joining tree built from a distance matrix computed using pseudohaploid base calls. Colours indicate management stocks. Dark green - OKH, light green - BCB, light blue ECWG, and dark blue - EGSB. Scale bar indicates pairwise distance.

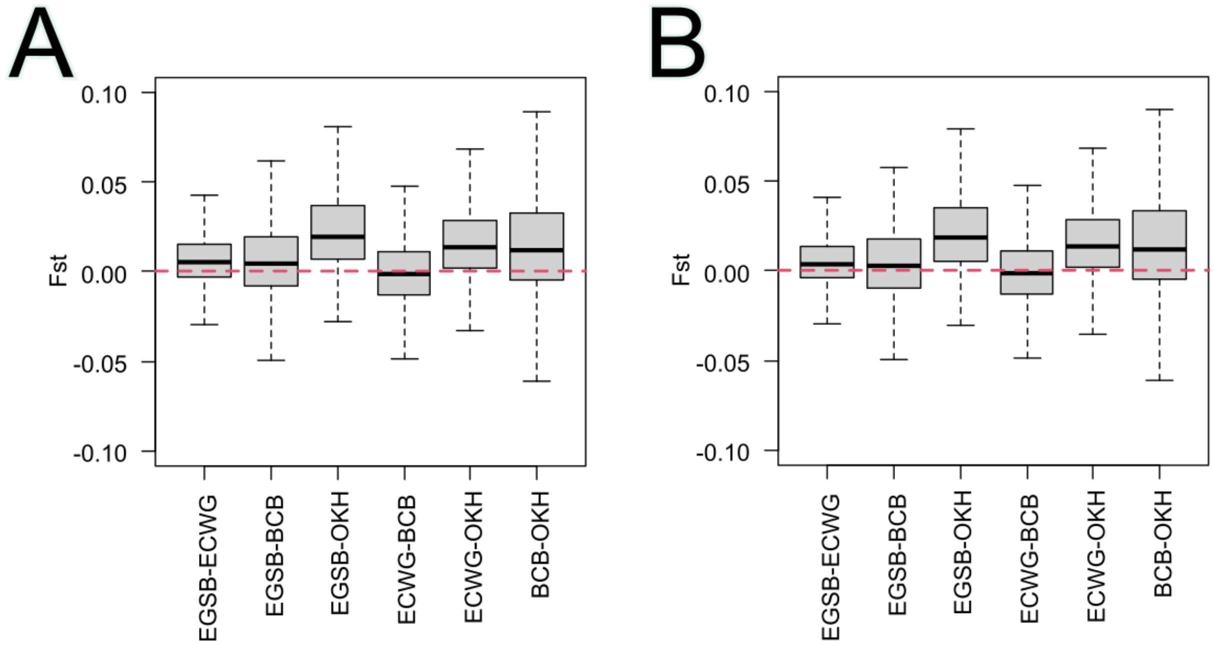


**Supplementary Figure S8: Evaluating the influence of coverage on Fst results. A)** Fst values computed from pseudohaploid base calls with no downsampling of EGSB. **B)** Fst values computed from pseudohaploid base calls with EGSB downsampled to ~2x.

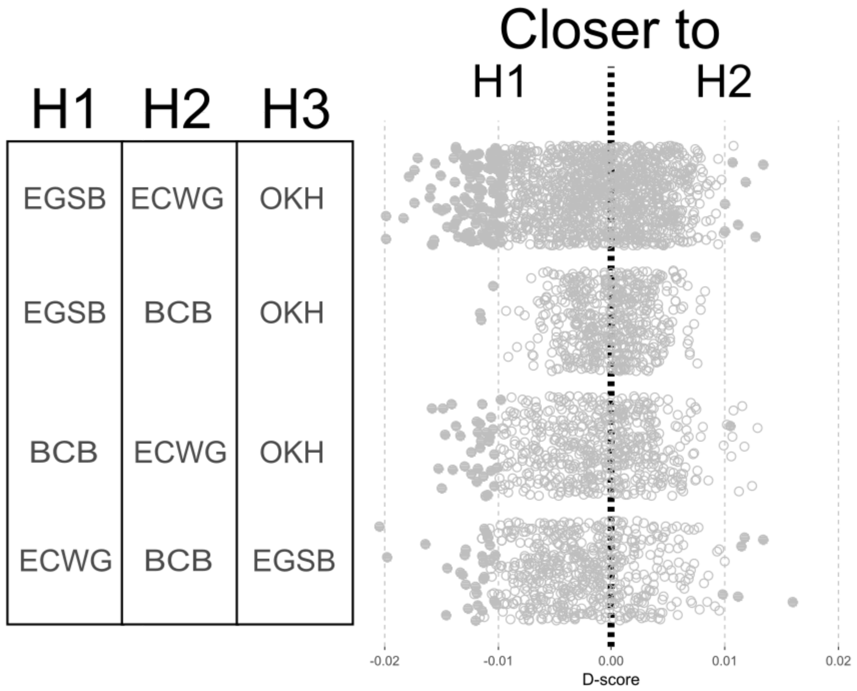


**Supplementary Figure S9:** D-statistics results calculated from a random base call approach. Filled circles show significant results (Z <|3|). Negative values show a closer relationship between H3 and H1 than H2. Positive results show a closer relationship between H3 and H2 than H1.

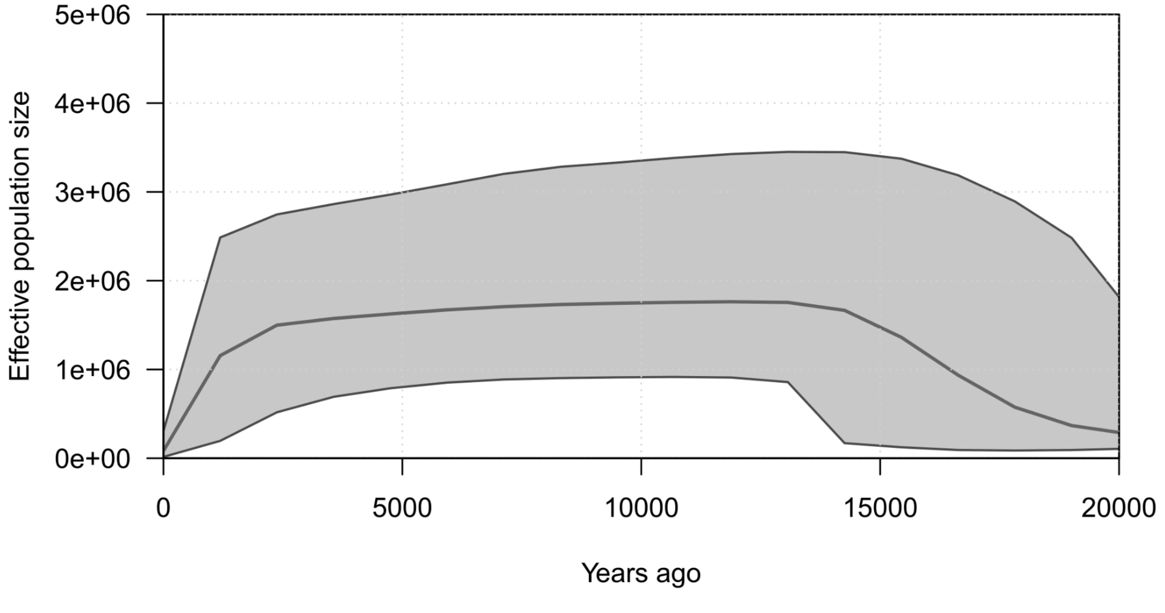


**Supplementary Figure S10:** Skyline plot built from the mitochondrial genomes and including the OHK individuals.

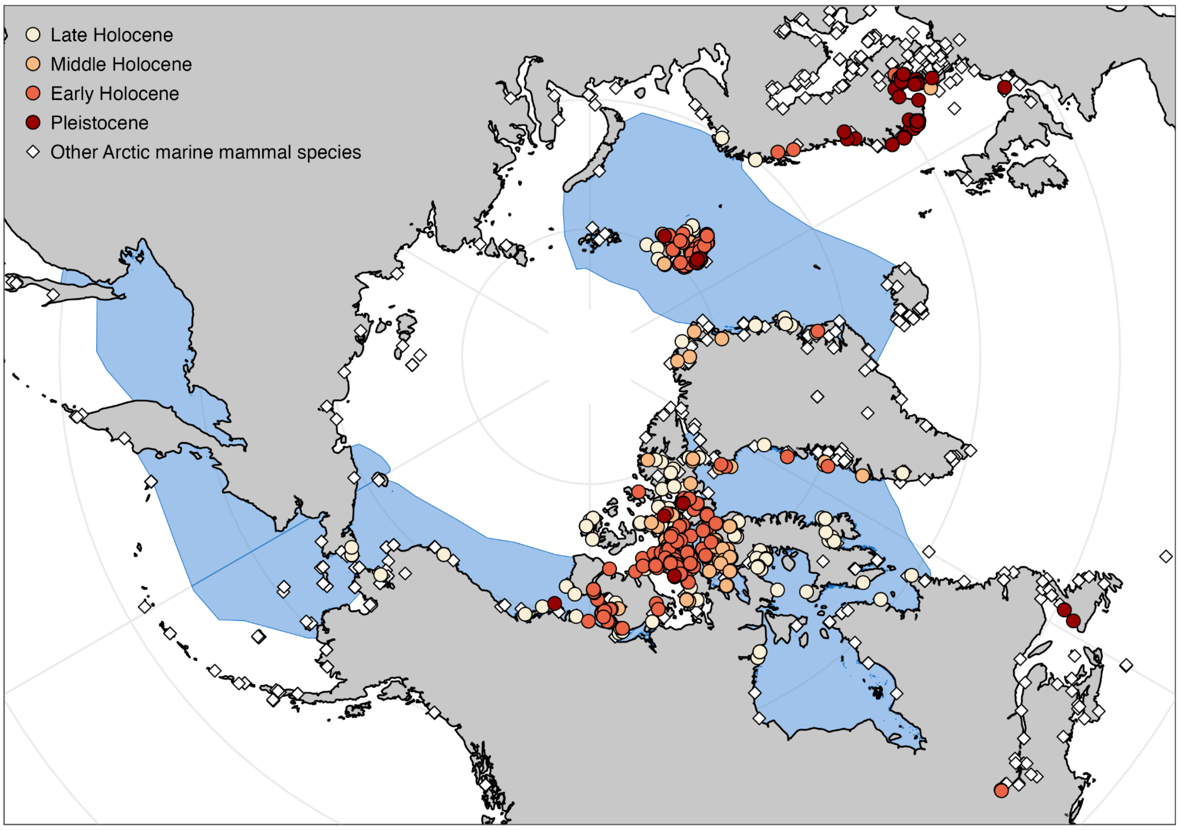


**Supplementary Figure S11 – Fossil record for bowhead whales and other Arctic marine mammals**. Location of bowhead whale (*n* = 1077) and other Arctic marine mammal (*n* = 2315) fossil remains in the Fossil Arctic Marine Mammal database (Freymueller et al. 2025b). Bowhead whale remains are shown in circles with colors denoting sample age. Other Arctic marine mammal species (11 species) are shown as diamonds. Blue areas denote current IUCN stock boundary polygons for bowhead whales.

**Supplementary tables:**

**Supplementary table S1: Sample information and genetic mapping statistics** - attached as spreadsheet

**Supplementary table S2: Population pairwise $F_{ST}$ values and $F_{ST}$ p-values based on the mitochondrial genomes.** The population $F_{ST}$ values are above the gray area and $F_{ST}$ p-values below. Bolded $F_{ST}$ value indicates significance (p<0.01).

| | **EGSB** | **ECWG** | **BCB** | **OKH** |
|---|---|---|---|---|
| **EGSB** | - | 0.10819 | 0.06953 | **0.20345** |
| **ECWG** | 0.03027+-0.0042 | - | 0.04908 | 0.09490 |
| **BCB** | 0.16113+-0.0117 | 0.14844+-0.0096 | - | 0.03413 |
| **OKH** | 0.00391+-0.0019 | 0.02344+-0.0048 | 0.29688+-0.0143 | - |

**Supplementary table S3: Mitochondrial diversity statistics** - n = number of samples, S = number of segregating sites, h = number of haplotypes

| Stock | n | *S* | *h* | Haplotype diversity | Nucleotide diversity ($\pi$) |
|---|---|---|---|---|---|
| EGSB | 15 | 139 | 9 | 0.933 (SD : 0.04) | 0.00292 (SD : 0.00017) |
| BCB | 7 | 104 | 7 | 1.00 (SD : 0.076) | 0.00255 (SD : 0.00068) |
| OKH | 11 | 99 | 4 | 0.745 (SD : 0.098) | 0.00207 (SD : 0.00061) |
| ECWG | 15 | 153 | 12 | 0.971 (SD : 0.033) | 0.00209 (SD : 0.00032) |
| All | 41 | 282 | 32 | 0.979 (SD : 0.009) | 0.00264 (SD : 0.0002) |

**Supplementary table S4: Genetic load values of the three inferred bowhead whale populations**. Genetic load values, calculated using SNPeff, were normalised by dividing by the number of homozygous synonymous mutations relative to the right whale outgroup. Realised load indicates the variant is in a homozygous state and masked load indicates a heterozygous state. LoF = Loss of function.

| **Individual ID** | **17-19** | **CGG_1_024320** | **CGG_1_024474** |
|---|---|---|---|
| **Population** | **EGSB** | **ECWG/BCB** | **OKH** |
| **Missense - masked** | 0.6437 | 0.5872 | 0.5931 |
| **Missense - realised** | 0.0810 | 0.0824 | 0.0913 |
| **LoF - masked** | 0.0320 | 0.0303 | 0.0280 |
| **LoF- realised** | 0.0036 | 0.0045 | 0.0048 |
